# READRetro: Natural Product Biosynthesis Planning with Retrieval-Augmented Dual-View Retrosynthesis

**DOI:** 10.1101/2023.03.21.533616

**Authors:** Seul Lee, Taein Kim, Min-Soo Choi, Yejin Kwak, Jeongbin Park, Sung Ju Hwang, Sang-Gyu Kim

**Affiliations:** Kim Jaechul Graduate School of AI, KAIST, Daejeon, 34141, Republic of Korea; Department of Biological Sciences, KAIST, Daejeon, 34141, Republic of Korea; Department of BioMedical Convergence Engineering, Pusan National University, Republic of Korea; School of Computing, KAIST, Daejeon, 34141, Republic of Korea

## Abstract

Elucidating the biosynthetic pathways of natural products has been a major focus of biochemistry and pharmacy. However, predicting the whole pathways from target molecules to metabolic building blocks remains a challenge. Here we propose READRetro as a practical bio-retrosynthesis tool for planning the biosynthetic pathways of natural products. READRetro effectively resolves the tradeoff between generalizability and memorability in bio-retrosynthesis by implementing two separate modules; each module is responsible for either generalizability or memorability. Specifically, READRetro utilizes a rule-based retriever for memorability and an ensemble of two dual-representation-based deep learning models for generalizability. Through extensive experiments, READRetro was demonstrated to outperform existing models by a large margin in terms of both generalizability and memorability. READRetro was also capable of predicting the known pathways of complex plant secondary metabolites such as monoterpene indole alkaloids, demonstrating its applicability in the real-world bio-retrosynthesis planning of natural products. A website (https://readretro.net) and open-source code have been provided for READRetro, a practical tool with state-of-the-art performance for natural product biosynthesis research.

## Introduction

Natural products are considered to be the most important source of molecules for a variety of applications, including dyes, fragrances, flavors, and small-molecule drugs^1^. In particular, 64.9% of small-molecule drugs approved by the Food and Drug Administration (FDA) since 1981 are natural products or their derivatives^2^. However, limited extraction from native organisms has been a major bottleneck in the study of native functions of pure compounds and their large-scale production^3^. Synthetic biology is considered to be a plausible solution for the efficient production of natural products using microorganisms as bio-based factories. Because metabolic engineering for producing natural products is based on the knowledge of biosynthesis in organisms, it is necessary to uncover the biosynthetic pathways of native hosts^3,4^. Recently, multi-omics approaches have been applied for identifying the biosynthetic pathways of complex molecules, such as colchicine and vinblastine^5–8^. However, it remains difficult to reveal intermediate molecular steps based on the complexity of searching for intermediate steps in a vast chemical space^5,9^. Therefore, planning feasible molecular pathways is an important goal in biochemistry.

In the era of machine learning, deep-learning-based approaches have emerged to predict complete synthetic pathways from molecules of interest to basic compound sets. This process is called retrosynthesis^10–12^. Deep-learning-based retrosynthesis models typically have a two-level structure in which a single-step retrosynthesis model predicts the reactants of a product, and a multi-step model performs a tree search in the chemical space by executing the single-step model iteratively^13^. Such models have primarily been developed in the field of organic chemistry^11,12,14–16^. However, organic retrosynthesis models may not be applicable when predicting the biosynthetic pathways of natural products because natural product biosynthesis involves specialized reaction mechanisms^17^. Accordingly, several retrosynthesis models for natural products have been developed based on biochemical reactions (bio-retrosynthesis)^18–22^. For instance, BioNavi-NP^22^, a template-free model, achieved state-of-the-art performance for predicting natural product reactions.

The goal of bio-retrosynthesis is to discover the metabolic pathways constructed by enzymatic reactions^18^. Metabolic pathways in organisms are highly conserved in metabolic groups and diversified through further modifications and combinations^23^. Based on these characteristics, an ideal bio-retrosynthesis model is required to predict unknown pathways (i.e., generalizability) and retrieve known reactions in its predictions (i.e., memorability)^17,22^. However, there is a tradeoff between generalizability and memorability because of the bias-variance tradeoff^24^. Therefore, existing bio-retrosynthesis models have limitations in terms of predicting either novel pathways or pathways consisting of known reactions^17^. However, no attempt has been made to overcome this bias-variance tradeoff by combining two separate models; each model is responsible for either generalizability or memorability.

Template-free retrosynthesis models learn conversion rules from products to reactants based on molecular SMILES (simplified molecular input line entry system) and/or graph representations. Chemical structures are represented using line notation with atomic symbols and bonds in SMILES, and as nodes (atoms) and edges (bonds) in graph representation^25^. SMILES-based models consider retrosynthesis as a neural machine translation task, mostly using a transformer architecture, and have become the dominant model for single-step retrosynthesis based on the ability of a transformer to capture global information^10,26^. However, SMILES-based models are constrained in depicting molecular information, which can limit their validity and diversity, negatively impacting generalizability^27^. In contrast, graph representations are considered to accurately reflect molecular characteristics, which has led to great advances in the field of cheminformatics, such as molecular property prediction and molecule generation^25^. Based on the permutation invariance and subgraph interpretability, the application of graph representations to retrosynthesis has been fruitful^27–37^. Recently, several retrosynthesis models utilizing both SMILES and graph representations have achieved superior performance on USPTO-50k benchmark dataset, which demonstrates the multifaceted and complementary nature of these two molecular representations^29,31,32,34^.

In this study, we introduce retrieval-augmented dual-view retrosynthesis (READRetro) as a novel tool for planning natural product biosynthetic pathways. READRetro utilizes two dual-representation-based single-step retrosynthesis models to take several advantages from graph representation. By constructing an ensemble of two single-step models, our single-step retrosynthesis model achieved state-of-the-art top-k accuracy on a metabolite reaction test set^22^. Furthermore, we developed a multistep retrosynthesis planning model by combining a retriever with the single-step retrosynthesis model and applying the Retro* algorithm^38^. READRetro outperformed baseline models in terms of all three metrics of biosynthetic pathway planning performance on two datasets: BioChem + USPTO_NPL for evaluating memorability and BioChem + USPTO_NPL (clean) for evaluating generalizability. An ablation study revealed that the dual representation of molecules improves the top-k accuracy of single-step retrosynthesis. We also provide a user-friendly web page and open-source code so that anyone can utilize READRetro for research purposes. With state-of-the-art performance in both generalizability and memorability based on the synergy of a dual-view deep learning-based model and a retriever, READRetro achieves a remarkable advance in the field of natural product biosynthetic pathway planning.

## Results

To evalute the performance of our model, READRetro was compared to baseline models (BioNavi-NP and RetroPathRL) trained on two datasets: BioChem + USPTO_NPL^22^ for evaluating generalizability and BioChem + USPTO_NPL (clean) for evaluating memorability. We used the multi-step 368 test set and BioChem + USPTO_NPL dataset from BioNavi-NP^22^. Because most reactions (92.9%) in the multi-step test set were already present in the BioChem + USPTO_NPL training set, the performance of models trained on BioChem + USPTO_NPL was mostly related to memorability. To evaluate the generalizability, BioChem + USPTO_NPL (clean) was constructed by removing the reactions in the multi-step test set from the BioChem + USPTO_NPL training set (see the dataset description in the Methods section for additional details).

### Evaluation of the single-step retrosynthesis

The performance of a single-step retrosynthesis model is critical for accurate multi-step pathway planning because multi-step retrosynthesis is achieved by iterating single-step predictions and performing a tree search. In this study, we used two dual-representation-based single-step retrosynthesis models called Retroformer^34^ and Graph2SMILES^29^. Retroformer is a transformer^26^-based model that exploits the parallel merging of SMILES and graph information through an encoder-decoder architecture with local-global attention heads. On the other hand, Graph2SMILES serially utilizes graph and SMILES information by incorporating a graph neural network encoder that directly captures the graph topology in front of the transformer. Due to the complementary nature of the dual molecular representations, Retroformer and Graph2SMILES have become state-of-the-art single-step retrosynthesis models. Because these two models utilize dual representations in different ways, we implemented an ensemble of Retroformer and Graph2SMILES to achieve an enhanced dual-view molecular representation. This ensemble model merges the results outputted by each individual model and produces top-10 proposals based on softmax-normalized scores.

The performance of the established ensemble model for predicting single-step metabolic reactions was evaluated (Table 1). The ensemble of Retroformer^34^ and Graph2SMILES^29^ achieved a top-1 accuracy of 23.4% and a top-10 accuracy of 59.3% on BioChem + USPTO_NPL, outperforming BioNavi-NP^22^ (18.6% and 58.6%, respectively). For BioChem + USPTO_NPL (clean), the ensemble consistently outperformed BioNavi-NP (58.1% versus 50.3% top-10 accuracy), demonstrating its robustness to a reduction in training data. Although BioNavi-NP is also an ensemble of four same architecture models (Transformer)^22,26^, it utilizes only SMILES without graph information, which limits its complementarity. The proposed ensemble of two dual-representation models exhibited superior prediction accuracy, indicating its excellent generalizability. We further evaluated the individual contributions of Retroformer and Graph2SMILES to the ensemble. The individual models underperformed compared to the ensemble, confirming the enhanced generalizability of the ensemble. We also evaluated the performance of Retroformer without atom mapping to examine the effects of graph representation. The Retroformer without atom mapping exhibited poor performance (52.9% and 52.0% top-10 accuracy on BioChem + USPTO_NPL and BioChem + USPTO_NPL (clean), respectively) compared to Retroformer (58.1% and 57.1% top-10 accuracy on BioChem + USPTO_NPL and BioChem + USPTO_NPL (clean), respectively). These results show that our ensemble model with an enhanced dual view achieves state-of-the-art predictive performance and generalizability. Therefore, we introduced this ensemble model into the multi-step planning framework described in the following section.

**Table 1.**
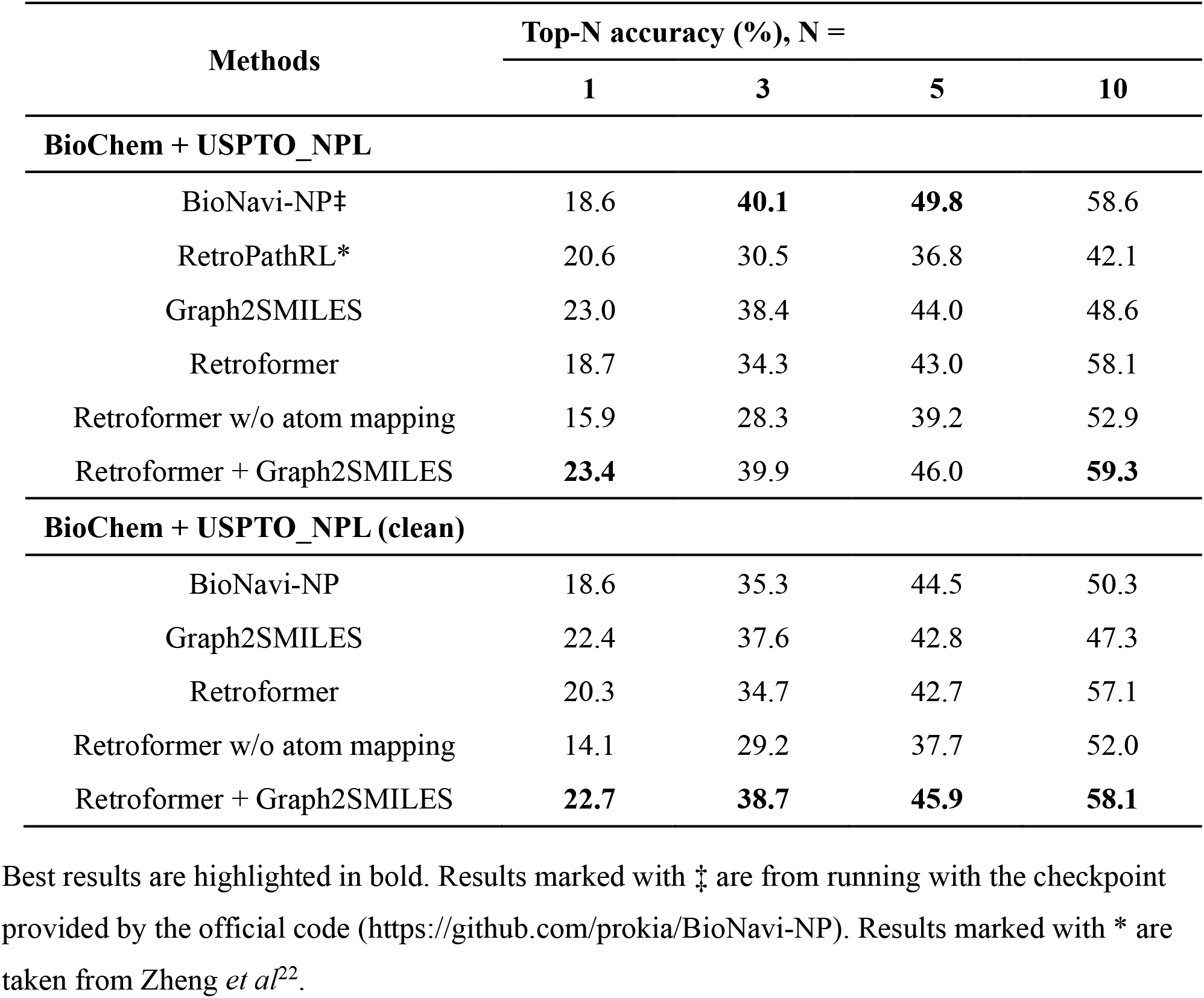
Performance of single-step retrosynthesis models.

### Evaluation of the multi-step retrosynthesis

Multi-step retrosynthesis planning is the ultimate goal of a retrosynthesis model. We constructed a multi-step retrosynthesis model called READRetro by applying the single-step retrosynthesis model (an ensemble of Retroformer and Graph2SMILES) and a retriever in the Retro* algorithm^38^. Notably, our retriever is a novel rule-based module developed to retrieve reactant molecules directly from a reaction database using product molecules as queries. Existing retrieval-based models retrieve reactant molecules from a predefined molecule set using deep-learning-based methods^39–41^. READRetro consists of a rule-based retriever and a deep-learning-based model, so it can fully leverage both the memorability of the retriever and generalizability of the deep learning model. Following BioNavi-NP^22^, the multi-step planning performance of the model was evaluated on 368 biosynthetic pathways (the test set) using three metrics: 1) success rate, the ability to plan routes from the target molecule to one of the predefined building blocks, 2) hit rate of building blocks, the ability to suggest routes that reach the exact building blocks used in the ground truth pathways, and 3) hit rate of pathways, the ability to plan synthetic routes that exactly match the ground-truth pathways. Intuitively, the success rate represents the scalability of a model, whereas the hit rate of building blocks and the hit rate of pathways indicate the validity of the pathways generated by a model. Because both scalability and validity are essential for retrosynthesis planning, a promising model should exhibit balanced performance across all metrics. The number of expansions and maximum number of iterations were set to 10 and 20, respectively. The maximum number of iterations in BioNavi-NP was set to 100, which is the default setting. The top-10 routes suggested by each model were used to evaluate the performance.

We first investigated the memorability of the retrosynthesis models by training them on BioChem + USPTO_NPL. As shown in Table 2, READRetro achieved a success rate of 93.2%, hit rate of building blocks of 87.2%, and hit rate of pathways of 66.6%, outperforming BioNavi-NP^22^ and RetroPathRL^19^ on all three metrics. In particular, a large improvement was observed in terms of validity according to the hit rates of building blocks and pathways. To analyze the contribution of each element of READRetro, we constructed multi-step models by applying each element: retriever, Retroformer and Graph2SMILES, to the Retro* algorithm^38^. As expected, READRetro outperformed all of these individual models, highlighting the synergistic effect between its components. Graph2SMILES showed a higher hit rate of pathways (47.0%) and a lower success rate (78.8%) compared to Retroformer (39.7% and 90.5%, respectively), which was consistent with its high top-1 accuracy and the low top-10 accuracy in the single-step evaluation (Table 1). In addition, the hit rates of building blocks and pathways of Retriever without deep learning models (74.7% and 69.0%, respectively) were much higher than those of the baselines (BioNavi-NP and RetroPathRL), indicating that the retriever contributes significantly to READRetro’s memorability. The decrease in the performance of READRetro when the retriever was removed reaffirms the impact of a retriever on the model’s memorability. Overall, based on the synergistic effect of the retriever and dual-representation-based deep learning models, READRetro achieved state-of-the-art performance for memorizing known biosynthetic pathways and utilizing them for multi-step pathway prediction.

**Table 2.** Performance of multi-step retrosynthesis models.

| Methods | Success rate (%) | Hit rate of building blocks (%) | Hit rate of pathways (%) |
| --- | --- | --- | --- |
| **BioChem + USPTO_NPL** |  |  |  |
| RetroPathRL* | 52.7 | 4.8 | 3.8 |
| BioNavi-NP* | 90.2 | 56.0 | 24.7 |
| Retriever | 84.2 | 74.7 | **69.0** |
| Retroformer | 90.5 | 81.0 | 39.7 |
| Graph2SMILES | 78.8 | 67.7 | 47.0 |
| READRetro w/o retriever | 90.5 | 79.3 | 39.9 |
| READRetro | **93.2** | **87.2** | 66.6 |
| **BioChem + USPTO_NPL (clean)** |  |  |  |
| BioNavi-NP | 73.4 | 37.8 | 4.9 |
| Retroformer | 84.8 | 72.0 | 25.0 |
| Graph2SMILES | 68.2 | 51.3 | 16.0 |
| READRetro | **85.3** | **72.8** | **26.6** |

Best results are highlighted in bold. Results marked with \* are taken from Zheng *et al*<sup>22</sup>.

To further investigate the generalizability of READRetro, we utilized BioChem + USPTO_NPL (clean) as a training set. READRetro largely outperformed the BioNavi-NP on all of the three metrics. The hit rates of building blocks and pathways of READRetro (72.8% and 26.6%, respectively) were much higher than those of BioNavi-NP (37.8% and 4.9%, respectively). The hit rates of READRetro trained on BioChem + USPTO_NPL (clean) were even higher than those of BioNavi-NP trained on BioChem + USPTO_NPL that contained most of the reactions (92.9%) in the test pathways. Furthermore, compared to Retroformer, the performance of the READRetro increased the success rate (0.5%), hit rate of building blocks (0.8%), and hit rate of pathways (1.6%), demonstrating the complementarity between Retroformer and Graph2SMILES. This remarkable performance of READRetro indicates that the ensemble of Retroformer and Graph2SMILES exhibits powerful generalizability that is enhanced by molecular dual representations.

### READRetro performance by chemical classes

We investigated the performance of READRetro for various classes of natural products (Fig. 2). Consistent with the multi-step evaluation results, READRetro outperformed both Retroformer^34^ and Graph2SMILES^29^ on all classes. READRetro achieved a success rate of 99.2%, hit rate of building blocks of 99.2%, and hit rate of pathways of 85.7% for the mevalonic acid or methylerythritol phosphate (MVA/MEP). The other classes in descending order of scores were amino acids (AAs), cinnamic acid or shikimic acid (CA/SA), and acetic acid and malonic acid (AA/MA). READRetro achieved a hit rate of pathways of 69.9% for class AAs consisting of complex molecules such as non-ribosomal peptide, ribosomally synthesized and posttranslational modified peptides, and alkaloids derived from amino acids (Fig. 2). READRetro showed relatively poor hit rates of pathways in the AA/MA class, which are mainly composed of aliphatic compounds such as fatty acid derivatives (Fig. 2). We expected an improvement in READRetro performance when incorporating more reactions into the training set to learn the specific rules for aliphatic compound reactions. Interestingly, although Retroformer and Graph2SMILES used both SMILES and graph representations, the two models exhibited distinct performances depending on the chemical category. Retroformer showed a higher hit rate of pathways than Graph2SMILES on the CA/SA and Others classes, and Graph2SMILES performed better on the AAs and MVA/MEP classes.

**Fig. 1:**
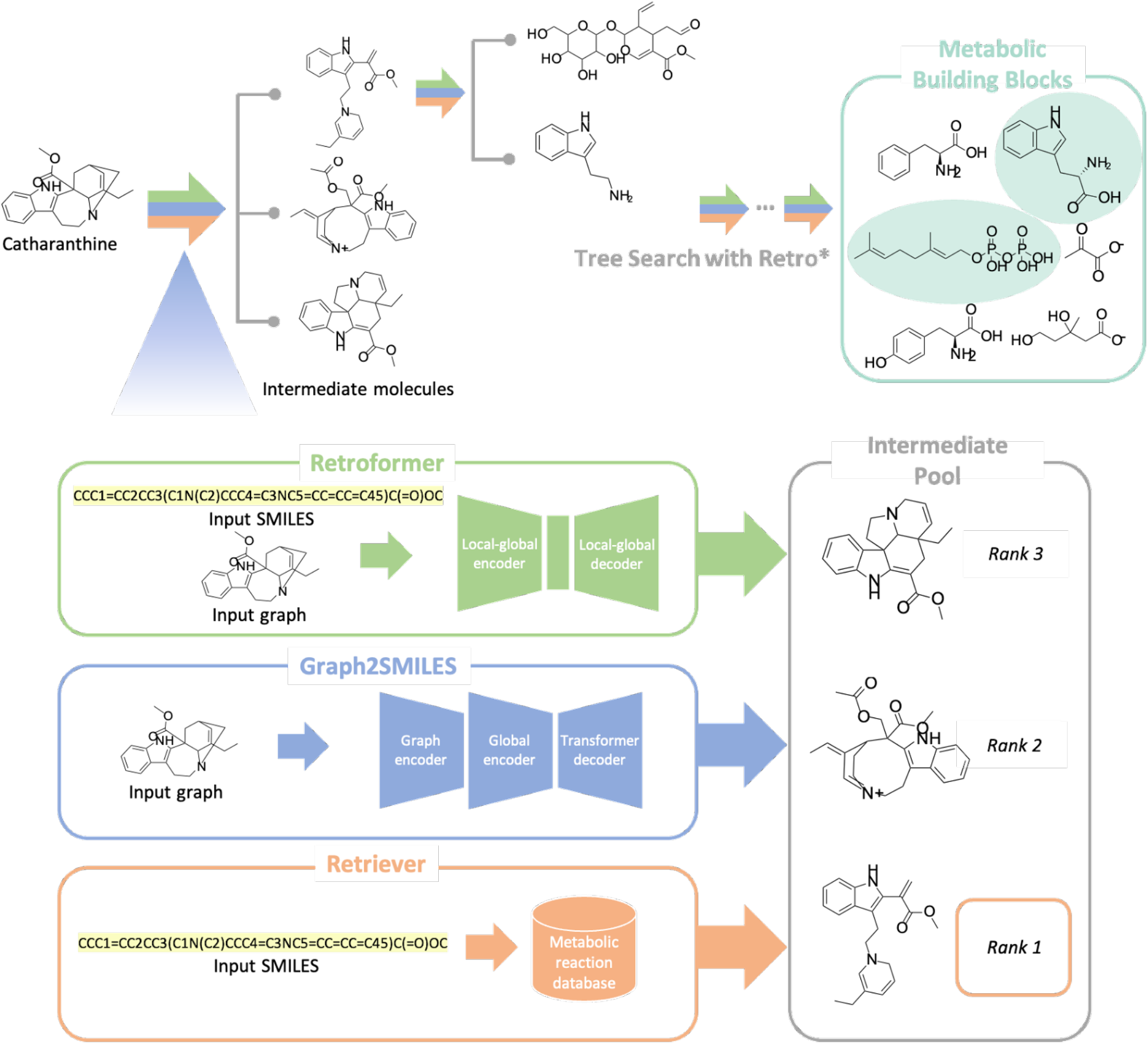
The overview of READRetro. READRetro is a bio-retrosynthesis model for planning the biosynthetic pathway of natural products. READRetro plans the biosynthetic pathway from target molecules to metabolic building blocks with an ensemble of deep learning-based chemical reaction prediction models, Retroformer and Graph2SMILES, and a reaction retriever. READRetro gives the highest score to the retrieved reactants to encourage the inclusion of prior knowledge from a large reaction database.

**Fig. 2:**
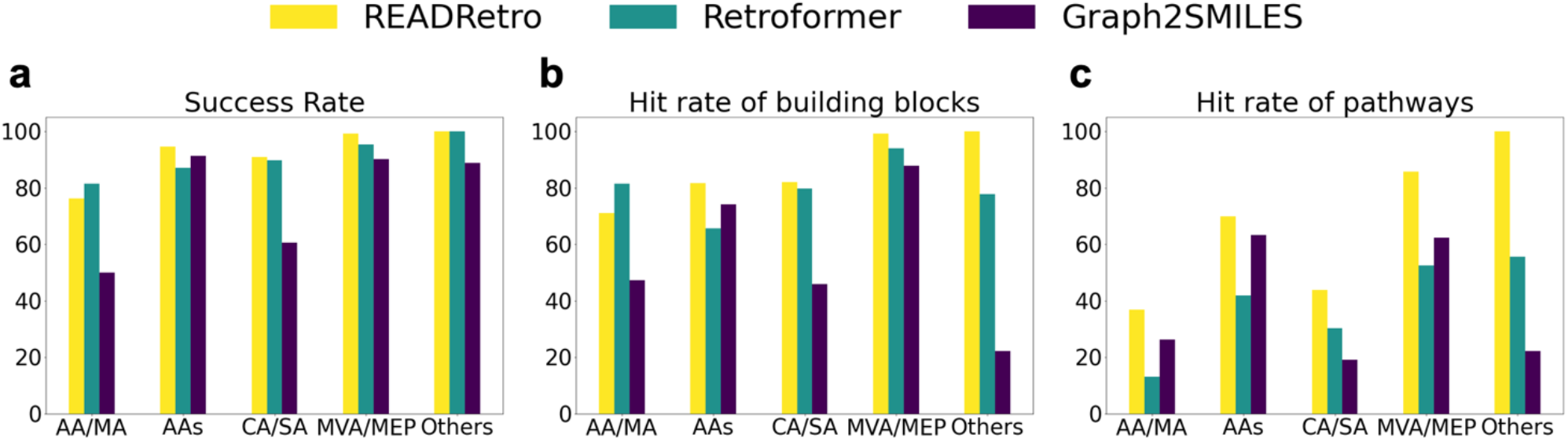
Performance of READRetro on multi-step test set by chemical classes. The success rates, the hit rates of building blocks, and the hit rates of pathways calculated by NP categories of **a** READRetro, **b** Retroformer, and **c** Graph2SMILES. The models were trained on BioChem + USPTO_NPL and evaluated on the 368 biosynthetic pathways of the multi-step test set. AA/MA, acetic acid and malonic acid; AAs, amino acids; CA/SA, cinnamic acid and shikimic acid; MVA/MEP, mevalonic acid and methylerythritol phosphate.

### Case study using READRetro

The main goal of a bio-retrosynthesis model is to aid in biological studies by predicting feasible synthetic pathways, which is often the first step in revealing complete synthetic pathways of natural products. To examine whether READRetro predicts pathways in accordance with previously established biosynthetic rules, we performed a case study on complex natural products. As target molecules become larger and more complex, both empirical retrosynthetic analysis and computer-aided retrosynthesis models rapidly lose accuracy. Monoterpene indole alkaloids (MIAs) are one of the structurally complex chemical families, and MIA’s biosynthetic pathways have been discovered by comprehensively integrating biochemical, genetic, and transcriptomic analyses^42–44^.

We tested whether READRetro and BioNavi-NP could reliably predict the biosynthetic pathways of two MIA compounds: tabersonine and catharanthine. In the biosynthetic pathways of MIAs, two building blocks called geranyl pyrophosphate (GPP) and tryptophan are metabolized separately and combined to form a key intermediate called strictosidine^43^. BioNavi-NP was unable to generate feasible synthetic pathways for tabersonine (Supplementary Fig. 2) or catharanthine (Supplementary Fig. 3). However, READRetro successfully predicted the complete biosynthetic pathways of tabersonine and catharanthine, including intermediate strictosidine and the building block tryptophan (Fig. 3a). The predicted synthetic pathways of tabersonine and catharanthine consisted of one generated final reaction (Fig. 3a, red box) and several database-retrieved steps (Fig. 3b, green box), highlighting the synergistic effect of the retriever and the deep-learning-based single-step model in READRetro. In addition, READRetro accurately predicted the biosynthetic pathways of other indole alkaloids, including elymoclavine (indole alkaloid; Fig. 3b) and 19E-geissoschizine (MIA; Supplementary Fig. 4). While BioNavi-NP generated irrelevant building blocks and intermediates for both compounds (Supplementary Figs. 5, 6), READRetro generated one final reaction (Fig. 3b, red box) and retrieved subsequent reactions (Fig. 3b, green box) to complete the retrosynthesis towards tryptophan. READRetro retrieved the correct complex synthetic pathways for harmine and baeocystin from the database (Supplementary Fig. 4). These results imply that READRetro can extract key components from complex natural products and predict their building blocks.

**Fig. 3:**
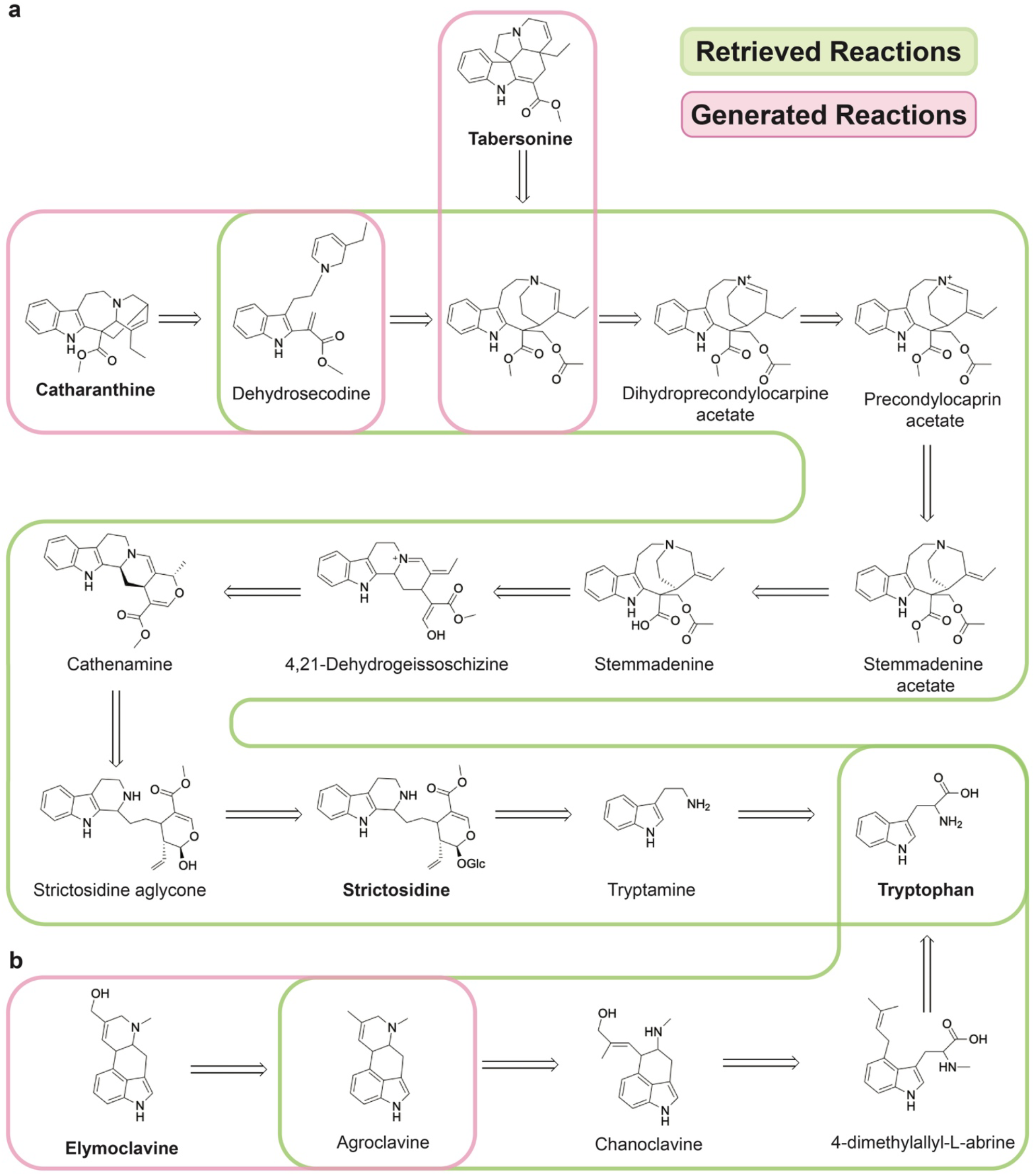
Case examples of READRetro. READRedtro predicts the correct biosynthetic pathways of **a** two monoterpene indole alkaloids: tabersonine and catharanthine and **b** one indole alkaloid, elymoclavine. Green boxes indicate retrieved reactions by the retriever and red boxes indicate generated reactions by the dual-view ensemble model. The maximum number of iterations was set to 100 as in BioNavi-NP.

We also conducted the retrosynthetic analyses of terpenoids, which are an important family of volatile and non-volatile compounds^45^. Monoterpenoids and sesquiterpenoids are synthesized from GPP and farnesyl pyrophosphate (FPP), respectively. Terpenoids are either directly synthesized from isoprenyl precursors by terpene synthases, or undergo further modifications to achieve enormous chemical diversity^46^. We tested several terpenoids for retrosynthetic predictions using READRetro. READRetro successfully retrieved known synthetic pathways (Supplementary Fig. 7, green box) for monoterpenoids and generated GPP as a building block when the reaction was not included in the dataset (Supplementary Fig. 7, red box). Likewise, the biosynthetic pathways of sesquiterpenoids towards FPP were also retrieved and/or generated by READRetro (Supplementary Fig. 8). Notably, three reactions were generated for the retrosynthesis of albicolide to reach the known pathway.

### Web implementation

READRetro was implemented on a web page (https://readretro.net) to provide it to the public a useful and practical tool. As shown in Fig. 4a, users can run READRetro by inputting the SMILES of target molecules, which can be obtained from web-based databases such as PubChem^47^. Users can change the following options: the number of generated pathways to display (default is 10), the maximum number of iterations (default is 100), the beam size (default is 10), the ON and OFF of the retriever (default is ON), and the building blocks (default is the 40 blocks depicted in Supplementary Fig. 9). Users can also change the set of building blocks in the “Building Blocks” tab to plan biosynthetic pathways reaching specific building blocks. Particular intermediates can be added to the building block set, which is effective for predicting pathways including intermediates. Users can also modify the retrieval database in the ‘Retrieval DB’ tab to apply more prior knowledge reactions or remove reactions for alternative pathway planning. Predicted pathways are shown in either the forward or reverse directions and in descending order of rank (Fig. 4b). In the predicted pathways, molecules found in the MetaNetX database^48^ are indicated by a blue background, and retrieved reactions are indicated by blue arrows. Predicted pathways can be filtered by intermediate/building blocks because feasible candidate pathways are typically determined by checking whether a pathway contains a specific intermediate and/or a building block. We highly recommend using the provided open-source code for those who wish to use READRetro intensively, or optimize our model for their own purposes.

**Fig. 4:**
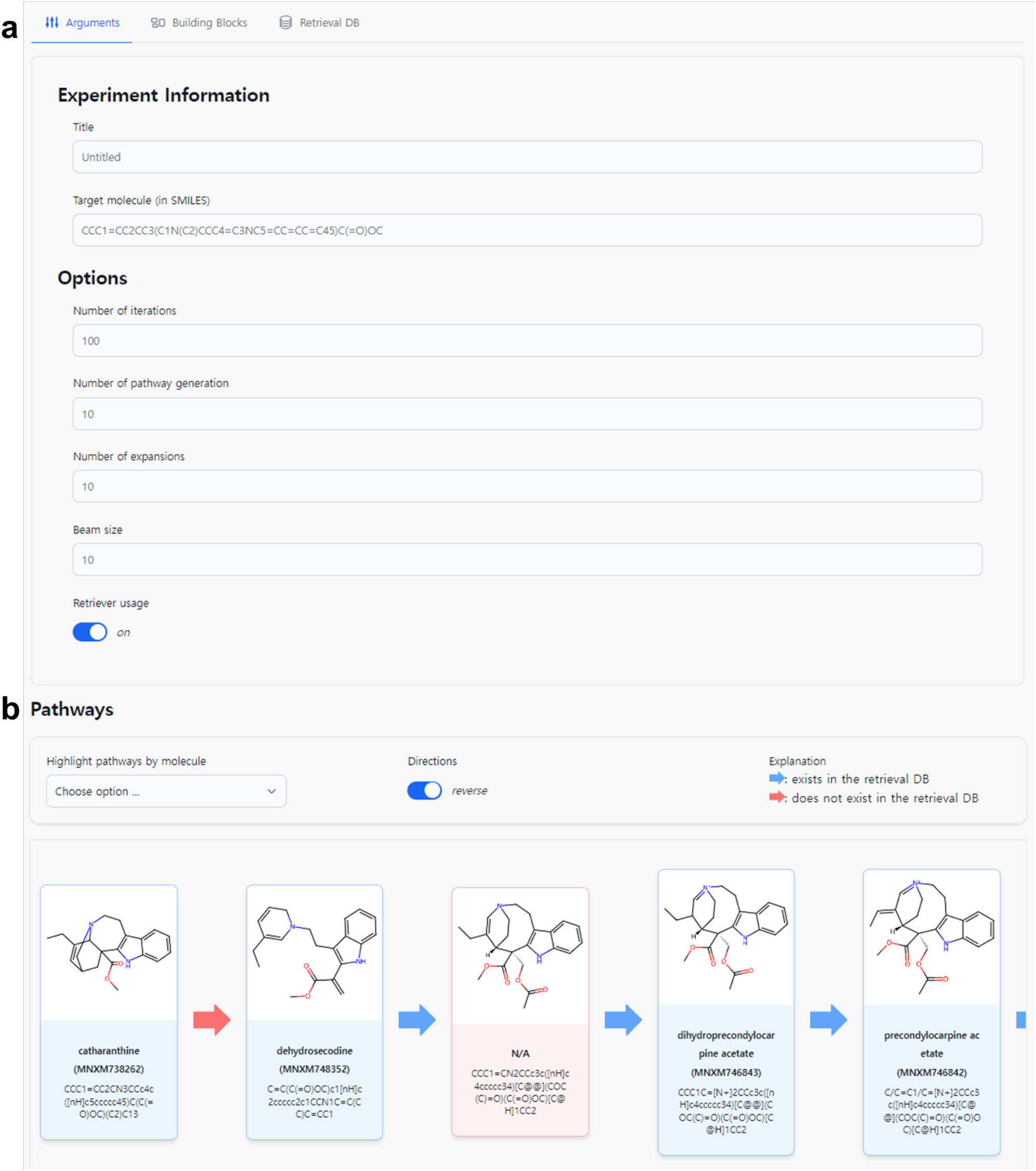
A web tool for READRetro. **a** Interface of web tool for READRetro **b** The results of planning biosynthetic pathway of catharanthine by READRetro web tool with default options. The intermediates found in MetaNetX^48^ database are indicated by blue background, and the blue arrows indicate the retrieved reactions.

## Discussion

In this work, we developed READRetro as a comprehensive model that can greatly facilitate biosynthetic pathway research on natural products. We pointed out that both generalizability and memorability are essential for predicting the conserved and diversified biosynthetic pathways of natural products. Existing retrosynthesis models either focus on only one of these two abilities or fail to overcome the tradeoff between the two abilities. To assess both generalizability and memorability, we proposed to evaluate multi-step bio-retrosynthesis models on two datasets: BioChem + USPTO_NPL and BioChem + USPTO_NPL (clean). Our evaluation protocol also included previously presented evaluation experiments^22^ for a fair and consistent comparisons, excluding arbitrary selection.

To overcome the tradeoff between generalizability and memorability, we proposed integrating two separate modules responsible for each ability. For generalizability, READRetro adopted two dual-representation-based deep learning models: Retroformer^34^ and Graph2SMILES^29^. Although dual-representation-based single-step retrosynthesis models have achieved state-of-the-art accuracy on the benchmark dataset^29,31,32,34^, to the best of our knowledge, we are the first to apply them to multi-step bio-retrosynthesis planning. Because Retroformer requires atom mappings of reactions between products and reactants to extract graph-edit information accurately, we built an atom-mapped BioChem + USPTO_NPL dataset^22^ using RXNMapper^49^, which is a neural-network-based automatic atom-mapping algorithm. The atom-mapped dataset can be utilized for bio-retrosynthesis using graph-based models that require atom mapping.

To establish an enhanced dual-view model, we constructed an ensemble of Retroformer and Graph2SMILES. Single-step and multi-step experiments validated the effectiveness of this ensemble for retrosynthesis pathway prediction (Table 1,2). The differences in model performance across classes of natural products illustrates the complementary nature of the two dual-representation-based models and the importance of the ensembles (Fig. 2). The state-of-the-art performance of our ensemble model suggests that there is a room for improvement in retrosynthesis models through combination. The same ensemble method could be applied to a recently proposed end-to-end retrosynthesis model^50^ to enable better biosynthetic pathway planning.

Regarding memorability, READRetro adopted a retriever module inspired by the open-domain questions answering field in natural language processing^51^. Because the retriever works at a single step level, it can aid in the prediction of not only fully known multi-step pathways, but also pathways consisting of known and unknown reactions (Fig. 3 and Supplementary Figs. 4,7,8). The proposed retrieval-augmented multi-step planning framework assigns the highest scores to the retrieved reactants, encouraging the inclusion of prior knowledge from a large reaction databases. This strategy is particularly effective for biosynthetic pathway planning because the pathways of natural products are highly conserved and diversified from common intermediates^23^. Unlike previous deep learning models^19,22^ that only implicitly memorize reactions in the training set through training loss, we emphasize that our rule-based retriever module explicitly memorizes known reactions, which dramatically improves the performance of READRetro. We also provide the option of running READRetro without a retriever or after removing some reactions from the retrieval database to plan alternative pathways. Reactions can be added to the retrieval database to expand prior knowledge. Applying a retriever to a multi-step retrosynthesis model is highly recommended as a simple and powerful way for incorporating existing biochemical knowledge (Table 2).

Although READRetro is able to generate promising biosynthetic pathways, as shown in our case study, some challenges remain to be overcome. Under the Retro* algorithm^38^, the sum of the costs of each single-step reaction is used to order the predicted pathways. As a result, long pathways are naturally considered undesirable, and the model tends to favor shorter pathways. Prioritizing on short pathways does not always lead to the correct prediction of the biosynthetic pathways of natural products, even though short pathways are generally preferred in chemical synthesis^38^. For instance, MIA biosynthetic pathways involve synthetically omittable intermediates^43^. In addition, when a natural product is synthesized by merging multiple independent biosynthetic pathways, only one branch is selected for the same reason. To solve these problems, we suggest normalizing pathway scores by the number of reaction steps.

## Methods

### Dataset construction

We used an open-source metabolic and organic reaction dataset called BioChem, offered from a previous bio-retrosynthesis study by Zheng et al.^22^ BioChem was constructed from MetaCyc^52^, KEGG^53^, and MetaNetX^48^. The BioChem dataset was preprocessed by performing irrelevant agent removal, multi-product reaction decompostion, and CoA to * replacement, and we used the same training-testing-validation split as the aforementioned study. The BioChem training set was merged with the USPTO_NPL dataset to construct the final training set of BioChem + UPSTO_NPL^22^. To evaluate the multi-step planning performance of READRetro and the baseline models, 368 internal test cases were utilized as in the previous study. In addition, because most single-step reactions (92.9%) in the multi-step internal test set were included in the BioChem + USPTO_NPL training set, we also constructed BioChem + USPTO_NPL (clean) by removing the reactions contained in the BioChem + USPTO_NPL training set to evaluate performance under unknown reaction.

Many retrosynthesis prediction models, including Retroformer^34^, require atom-mapped training data between reactants and products. Because the reactions in BioChem + USPTO_NPL are not atom-mapped^22^, we utilized a neural network-based automated atom-mapping model called RXNMapper^49^ to generate an atom-mapped dataset. A total of 88,666 reactions were atom mapped from 93,958 reactions in the BioChem + USPTO_NPL training set (94.4%). Because RXNMapper^49^ canonicalizes reaction SMILES, reactions containing a “R” token in their SMILES string were removed. The reactions in the BioChem + USPTO_NPL (clean) training set were atom mapped to produce 88,167 reactions by the same procedure (Supplementary Fig. 1).

### Model

The goal of retrosynthesis prediction is to predict the precursors of a given target molecule by capturing underlying biochemical rules from a training database. We construct a single-step prediction model by combining three models: 1) a dual-representation-based sequence-to-sequence model, Retroformer^34^, 2) a dual-representation-based graph-to-sequence model, Graph2SMILES^29^, and 3) a rule based retriever^25^. Retroformer utilizes a global head identical to the self-attention head of a vanilla transformer^26^ and a local head that captures a graph topology to infer a reaction center. Graph2SMILES^29^ uses a graph neural network (GNN) encoder^54^ to obtain atom-level representations, which are then fed into a transformer^26^-based global attention encoder and decoder. All models were constructed, trained, and tested using the PyTorch^55^ library. We trained the Retroformer models for 1,600,000 iterations with the default settings specified in the official codebase (https://github.com/yuewan2/Retroformer). Graph2SMILES models were trained for 84,000 iterations on BioChem + USPTO_NPL and 72,000 iterations on BioChem + USPTO_NPL (clean) with the options described in Supplementary Data 1, which were optimized based on corresponding official training scripts (https://github.com/coleygroup/Graph2SMILES). The Retroformer and Graph2SMILES models each propose up to ten reactant candidates, which are then passed into a softmax function for normalization. The retriever memorizes the reactions in the training set and outputs the corresponding reactants for a given product. The reactant scores proposed by the retriever are set to maximum value of one. The beam sizes of the Retroformer and Graph2SMILES models were set to 10 and 200, respectively.

To perform multi-step retrosynthesis prediction, READRetro utilizes Retro*^38^, which is an A*- like tree search algorithm that iteratively expands the search tree using a single-step retrosynthesis model and neural cost function. The top-10 reactants and their scores from the single-step model are fed to Retro* in each expansion. Retro* constructs an AND-OR tree by considering molecules as OR nodes and reactions as AND nodes, and expands the tree according to the search bias of a pre-trained neural network that predicts synthesis costs. The reactant costs proposed by the retriever are set to a minimum cost of zero. The maximum number of search iterations was set to 20, and the 40 building blocks which are the precursors of most natural products were used following the previous work^22^. After tree expansion was complete, the top-10 routes in terms of total cost, which is defined as the sum of the single-step costs, were used to evaluate performance.

## Data availability

All dataset will be deposited after acceptance.

## Code availability

The web page of READRetro is available at https://readretro.net. All code generated during this study will be deposited after acceptance.

## Acknowledgments

This work was supported by grants from the National Research Foundation of Korea (NRF) grants (NRF-2018R1A5A7025409), the Rural Development Administration (PJ016993), the Post-AI program of KAIST, Institute of Information & communications Technology Planning & Evaluation (IITP) grant funded by the Korea government MSIT (No.2021-0-02068, Artificial Intelligence Innovation Hub and No.2019-0-00075, Artificial Intelligence Graduate School Program at KAIST), and the Engineering Research Center Program through the National Research Foundation of Korea (NRF) funded by the Korean government MSIT (NRF-2018R1A5A1059921). JP and YK were supported by Institute of Information & communications Technology Planning & Evaluation (IITP) grant funded by the Korea government MSIT (No.2020-2-01450, Artificial Intelligence Convergence Research Center, Pusan National University) and the National Research Foundation of Korea (NRF) grant funded by the Korea government MSIT (No. 2022R1F1A1076160).

## Author contributions

S.J.H and S.-G.K. conceived and supervised the project. S.L and T.K developed READRetro and evaluated the performance of READRetro compared to other retrosynthesis models. M.-S.C. performed the case study of READRetro. Y.K. and J.P developed the web version of READRetro. S.L, T.K, and M.-S.C. wrote the manuscript.

