## supplementary figure for "READRetro: Natural Product Biosynthesis Planning with Retrieval-Augmented Dual-View Retrosynthesis"

### Supplementary information

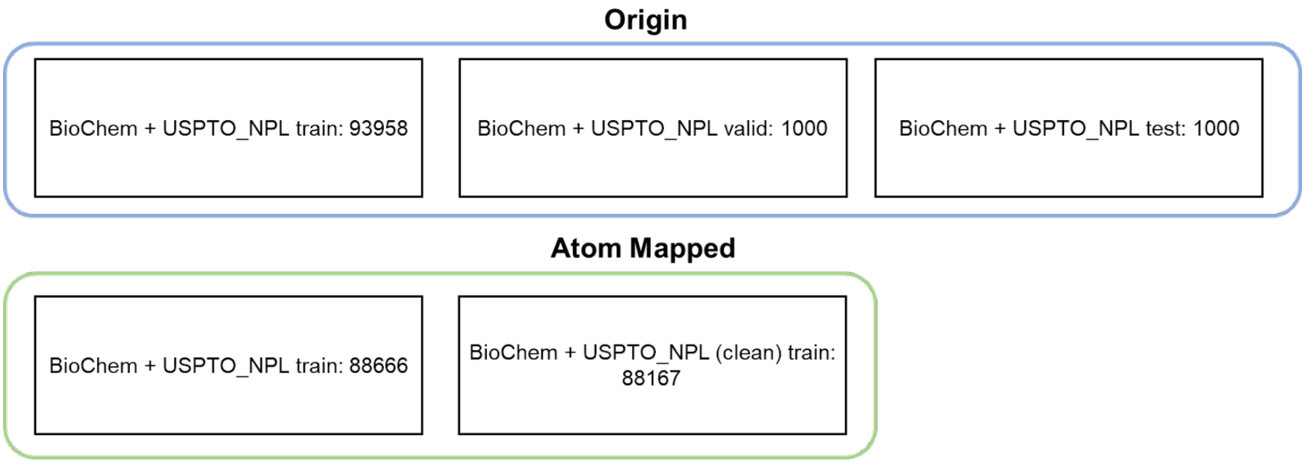


**Supplementary Figure 1. Dataset overview.** Datasets in Origin (blue box) are open sources offered by Zheng *et al*[22]. BioChem + USPTO_NPL train in Atom Mapped (green box) was atom-mapped from BioChem + USPTO_NPL train (blue box) by RXNMapper[49], and Atom Mapped BioChem + USPTO_NPL (clean) train (in green box) was constructed by removing all reactions in multi-step test set[22] from the Atom Mapped BioChem + USPTO_NPL train.

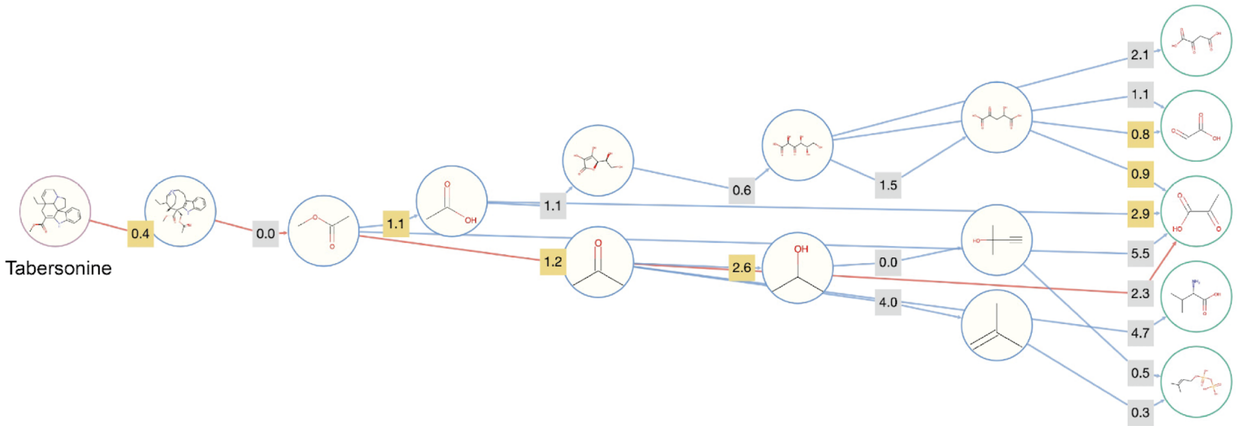


**Supplementary Figure 2. Predicted synthetic pathways of tabersonine by BioNavi-NP default options.**

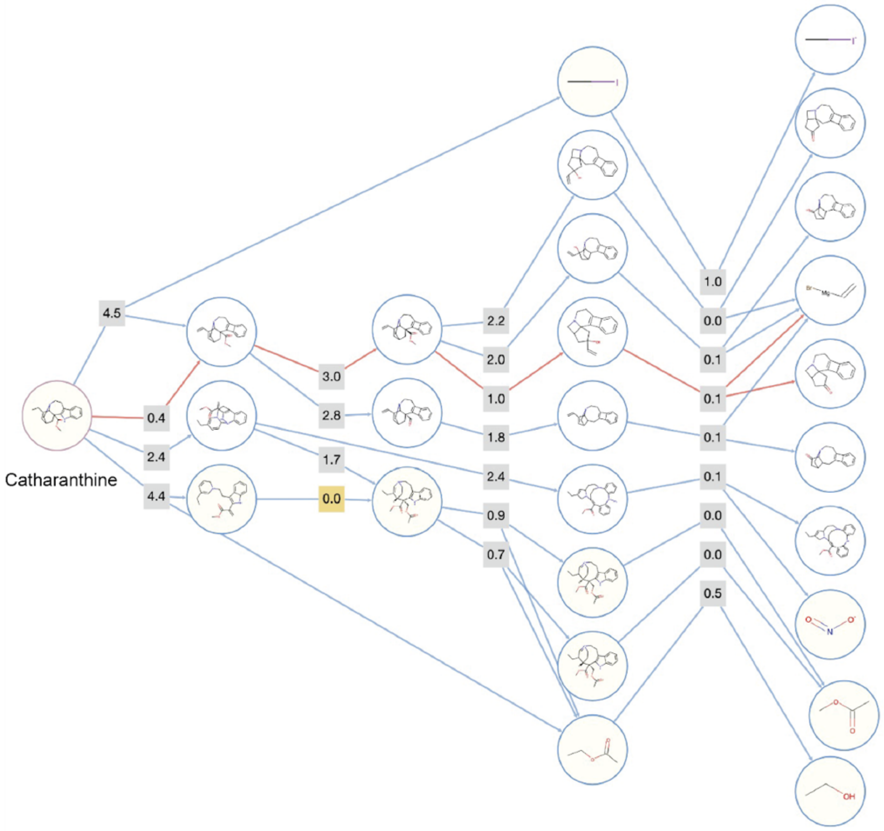


**Supplementary Figure 3. Predicted synthetic pathways of catharanthine by BioNavi-NP default options.**

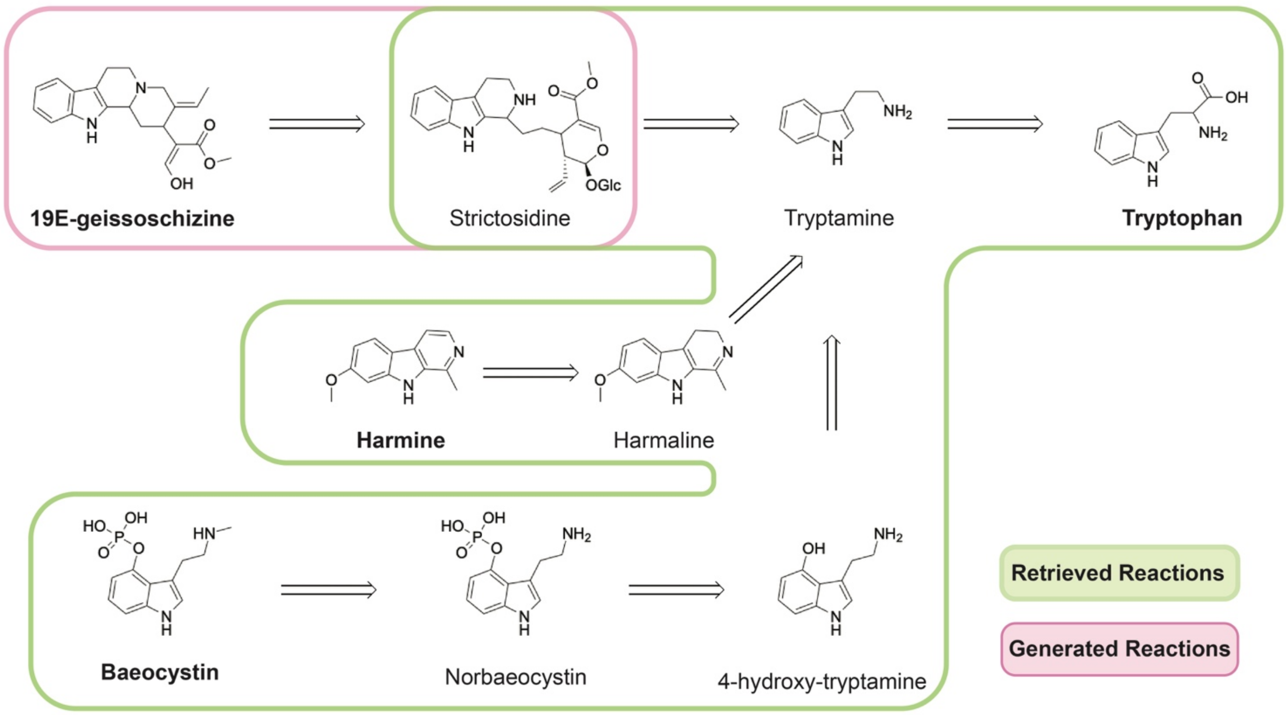


**Supplementary Figure 4. Predicted synthetic pathways of 19E-geissoschizine (MIA), baeocystin (IA), and harmine (IA) by READRetro.** Green box indicates retrieved reactions by the retriever, and red box indicates generated reactions by the dual-view ensemble model. The maximum number of iterations were set to 100 as in BioNavi-NP. MIA, monoterpene indole alkaloid; IA, indole alkaloid.

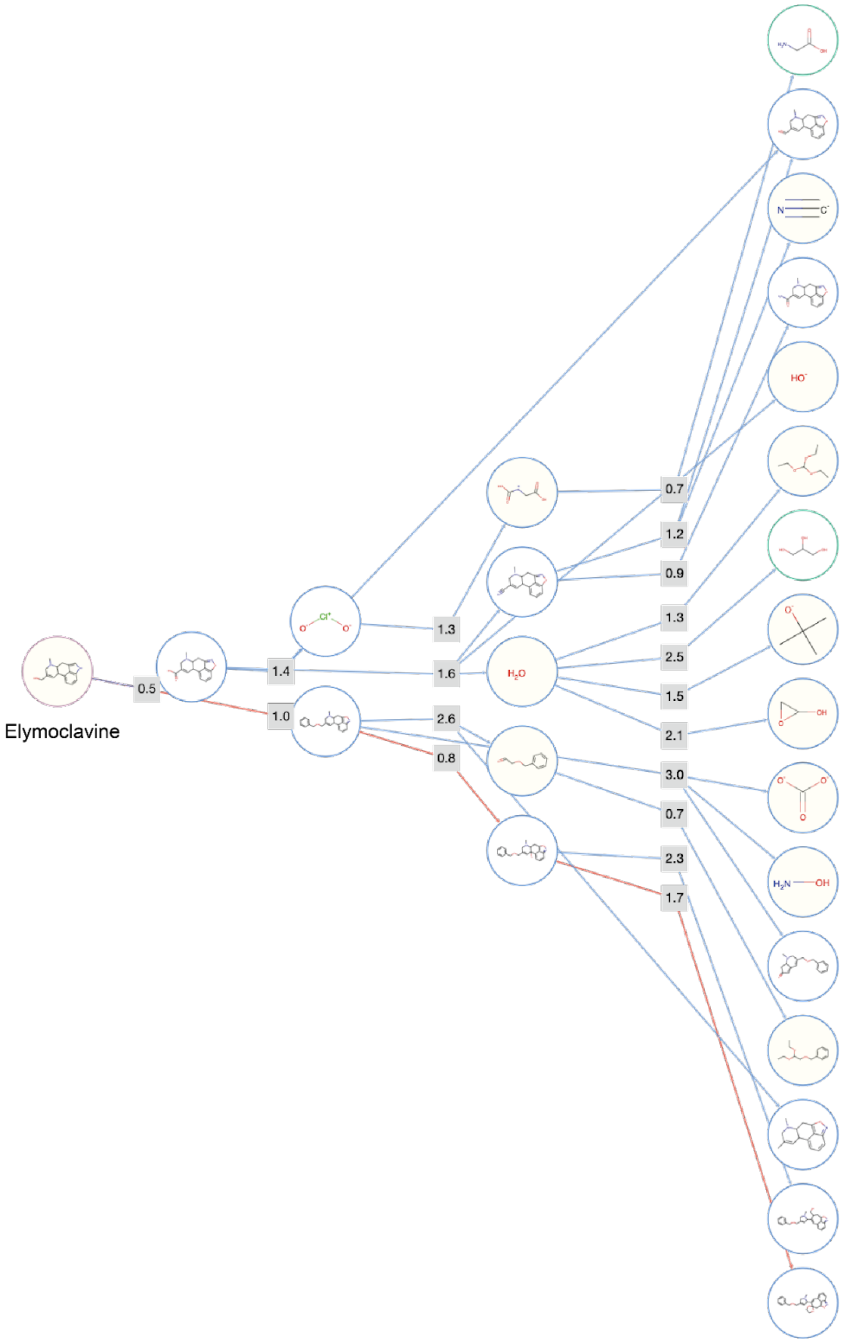


**Supplementary Figure 5. Predicted synthetic pathways of elymoclavine by BioNavi-NP default options.**

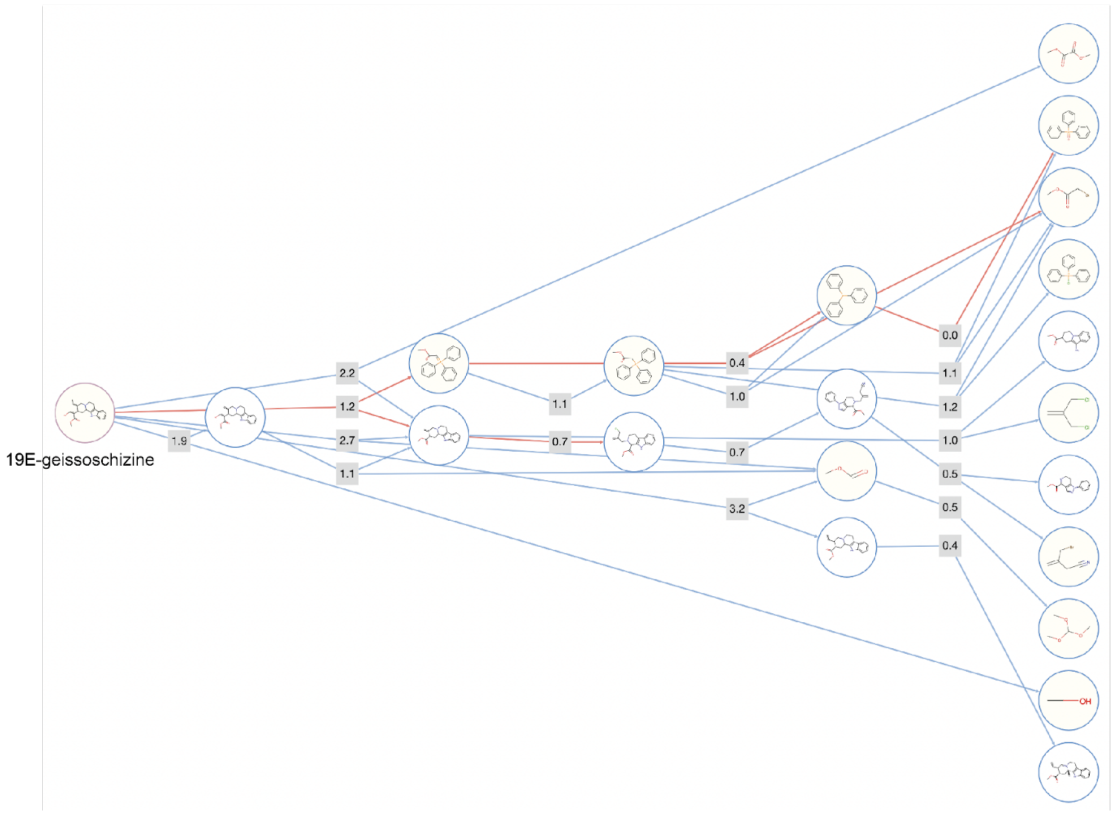


**Supplementary Figure 6. Predicted synthetic pathways of 19E-geissoschizine by BioNavi-NP default options.**

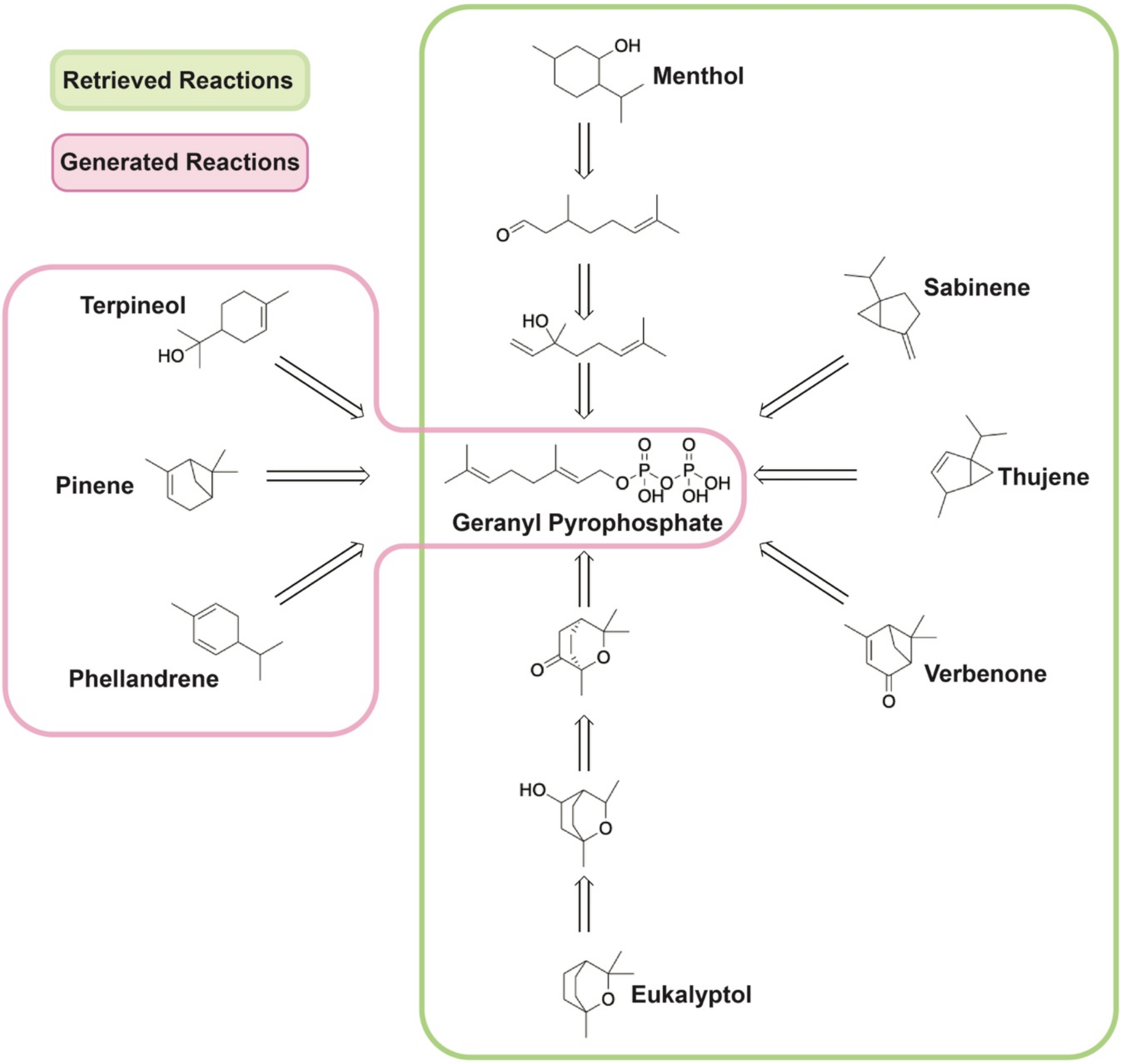


**Supplementary Figure 7. Predicted synthetic pathways of monoterpenoids by READRetro.** Green box indicates retrieved reactions by the retriever, and red box indicates generated reactions by the dual-view ensemble model. The maximum number of iterations were set to 100 as in BioNavi-NP.

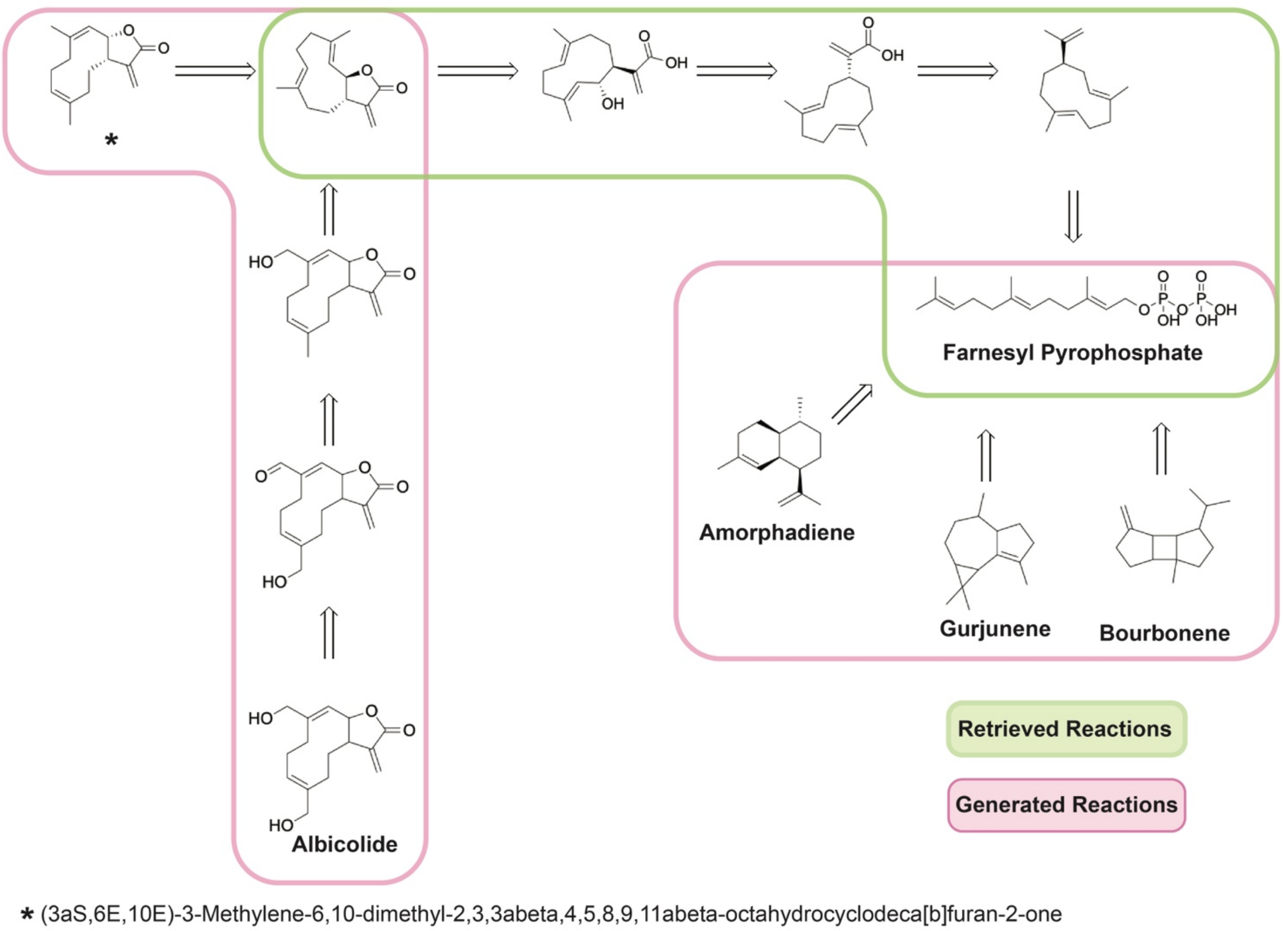


**Supplementary Figure 8. Predicted synthetic pathways of sesquiterpenoids by READRetro.** Green box indicates retrieved reactions by the retriever, and red boxes indicate generated reactions by the dual-view ensemble model. The maximum number of iterations were set to 100 as in BioNavi-NP.

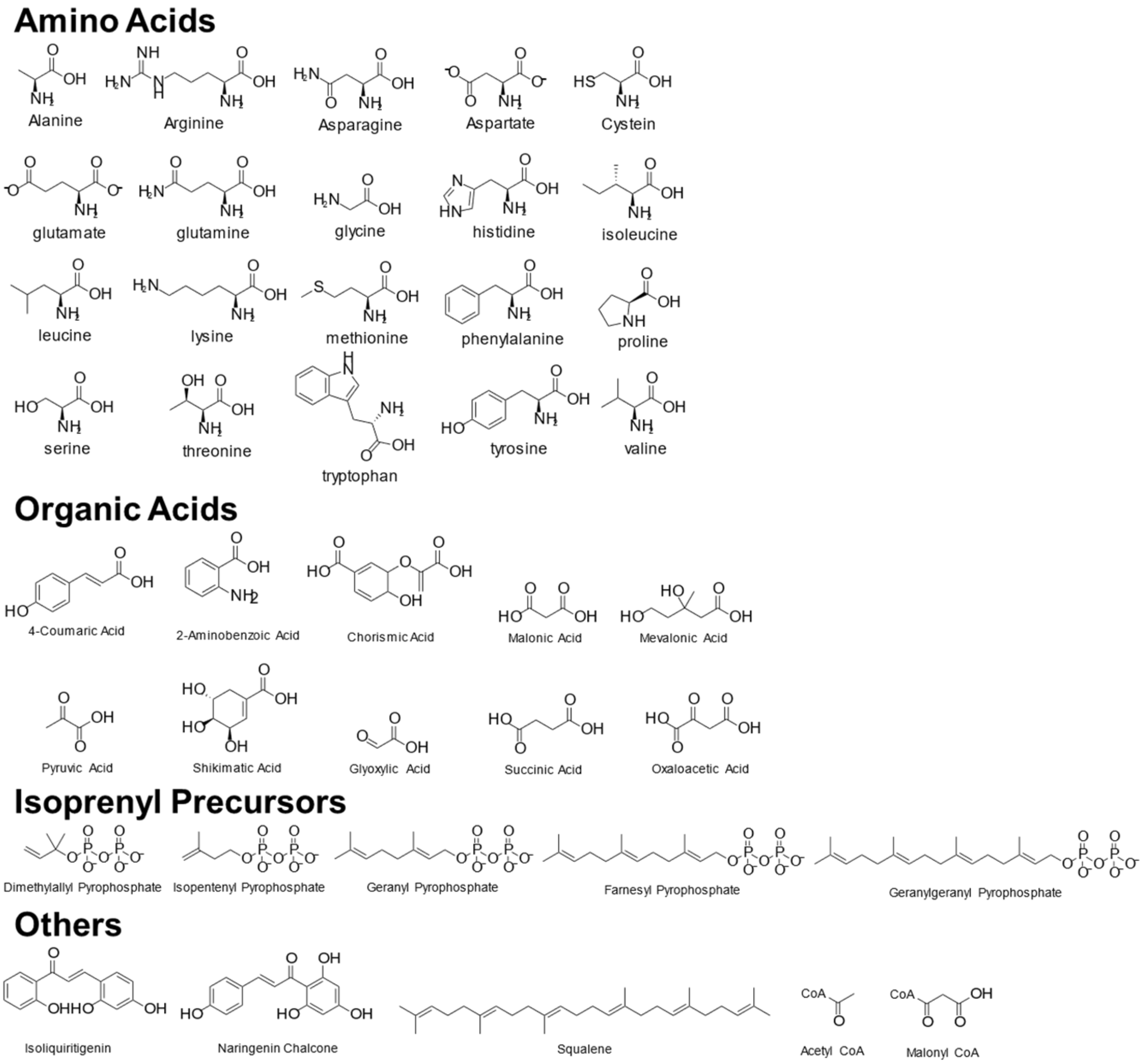


**Supplementary Figure 9. Building blocks of READRetro.**

### Supplementary Data 1. Implementation details of READRetro.

**a** Single-step implementation details of READRetro

To train Retroformer, we used the following command using the official code (https://github.com/yuewan2/Retroformer).

```
python retroformer/train.py \
  --encoder_num_layers 8 --decoder_num_layers 8 --heads 8 --max_step 2000000 \
  --batch_size_trn 4 --batch_size_val 4 --batch_size_token 4096 \
  --save_per_step 100000 --val_per_step 10000 --report_per_step 200 --device cuda \
  --known_class False --shared_vocab True --shared_encoder False \
  --data_dir data/$DATA_DIR --intermediate_dir intermediates/$INTERMEDIATE_DIR \
  --checkpoint_dir $SAVE_DIR
```

To train Graph2SMILES, we used the following command using the official code (https://github.com/coleygroup/Graph2SMILES).

```
python train.py \
  --model="g2s_series_rel" --data_name="$DATASET" --task="retrosynthesis" \
  --representation_end=smiles --train_bin="./preprocessed/$PREFIX/train_0.npz" \
  --valid_bin="./preprocessed/$PREFIX/val_0.npz" \
  --log_file="$PREFIX.train.$EXP_NO.log" \
  --vocab_file="./preprocessed/$PREFIX/vocab_$REPR_END.txt" \
  --save_dir="./checkpoints/$RESULT.$EXP_NO" --embed_size=256 \
  --mpn_type="dgcn" --encoder_num_layers="4" --encoder_hidden_size="256" \
  --encoder_norm="" --encoder_skip_connection="" \
  --encoder_positional_encoding="none" --encoder_emb_scale="sqrt" \
  --attn_enc_num_layers=8 --attn_enc_hidden_size=256 --attn_enc_heads=8 \
  --attn_enc_filter_size=2048 --rel_pos="emb_only" --rel_pos_buckets=11 \
  --decoder_num_layers=6 --decoder_hidden_size=256 --decoder_attn_heads=8 \
  --decoder_filter_size=2048 --dropout=0.3 --attn_dropout=0.3 \
  --max_relative_positions=4 --seed=42 --epoch=2000 --max_steps=2000000 \
  --warmup_steps=8000 --lr=2 --weight_decay=0.0 --clip_norm=20.0 \
  --batch_type="tokens" --train_batch_size=2048 --valid_batch_size=2048 \
  --predict_batch_size=2048 --accumulation_count=4 --num_workers=0 \
  --beam_size=10 --predict_min_len=1 --predict_max_len=512 --log_iter=100 \
  --eval_iter=2000 --save_iter=2000 --compute_graph_distance
```

The retriever was implemented by the following code.

```
from collections import defaultdict
from utils.ensemble import run_ensemble


class Retriever:
    def __init__(self, data):
        with open(data) as f:
            reactions = f.readlines()
        self.graph = defaultdict(list)
        for reaction in reactions:
            reactant, product = reaction.strip().split('>>')
            self.graph[product].append(reactant)

    def retrieve(self, product):
        if product in self.graph:
            return self.graph[product]


def run_retriever(product, retriever, model_retroformer, model_g2s,
                  args_retroformer, args_g2s,
                  vocab, vocab_tokens, device, expansion_topk):
    reactant_list = retriever.retrieve(product)
    if reactant_list is not None:
        res_dict = {'reactants': reactant_list}
        res_dict['scores'] = [1. for _ in res_dict['reactants']]
        res_dict['retrieved'] = [True for _ in res_dict['reactants']]
```

```
        _res_dict = run_ensemble(product, model_retroformer, model_g2s,
                                 args_retroformer, args_g2s,
                                 vocab, vocab_tokens, device, expansion_topk)
        res_dict['reactants'].extend(_res_dict['reactants'])
        res_dict['scores'].extend(_res_dict['scores'])
        res_dict['retrieved'].extend(_res_dict['retrieved'])
        res_dict['templates'] = [None for _ in res_dict['scores']]
    else:
        res_dict = run_ensemble(product, model_retroformer, model_g2s,
                                args_retroformer, args_g2s,
                                vocab, vocab_tokens, device, expansion_topk)
    return res_dict
```

The reactant scores proposed by the retriever are set to 1.

**b** Multi-step implementation details

We modified and adopt Retro*[38] to conduct a tree search with the proposed single-step model in READRetro. The options for running Retro* in the multi-step evaluation were the number of pathway generation of 10, the maximum number of iterations of 20, the number of expansions of 10, the beam size of 10. Implementation details could be found in the open-source code.

```
from retro_star.api import RSPlanner
import argparse

parser = argparse.ArgumentParser()
parser.add_argument('product',                  type=str)
parser.add_argument('-b', '--blocks',           type=str, default='data/building_block.csv')
parser.add_argument('-i', '--iterations',       type=int, default=20)
parser.add_argument('-e', '--exp_topk',         type=int, default=10)
parser.add_argument('-k', '--route_topk',       type=int, default=10)
parser.add_argument('-s', '--beam_size',        type=int, default=10)
parser.add_argument('-r', '--retrieval',        type=str, default='true', choices=['true',
'false'])
parser.add_argument('-d', '--retrieval_db', type=str,
default='data/train_canonicalized.txt')
parser.add_argument('-c', '--device',           type=str, default='cuda', choices=['cuda',
'cpu'])
args = parser.parse_args()

planner = RSPlanner(
    cuda=args.device=='cuda',
    iterations=args.iterations,
    expansion_topk=args.exp_topk,
    route_topk=args.route_topk,
    beam_size=args.beam_size,
    retrieval=args.retrieval=='true',
    retrieval_db=args.retrieval_db,
    starting_molecules=args.blocks
)
result = planner.plan(args.product)
if result is None:
    print('None')
else:
    for i, route in enumerate(result):
        print(f'{i} {route}')
```
